# Improved antibody-specific epitope prediction using AlphaFold and AbAdapt

**DOI:** 10.1101/2022.05.21.492907

**Authors:** Zichang Xu, Ana Davila, Jan Wiamowski, Shunsuke Teraguchi, Daron M. Standley

## Abstract

Antibodies recognize their cognate antigens with high affinity and specificity, but the prediction of binding sites on the antigen (epitope) corresponding to a specific antibody remains a challenging problem. To address this problem, we developed AbAdapt, a pipeline that integrates antibody and antigen structural modeling with rigid docking in order to derive antibody-antigen specific features for epitope prediction. In this study, we assess the impact of integrating the state-of-the-art protein modeling method AlphaFold with the AbAdapt pipeline and demonstrate significant improvement in the prediction of antibody-specific epitope residues.

## 1. Introduction

Highly specific antibody-antigen interactions are a defining feature of adaptive immune responses to pathogens or other sources of non-self molecules. This adaptive molecular recognition has been exploited to engineer antibodies for various purposes, including laboratory assays and highly specific protein therapeutics. Despite their widespread use, experimental identification of antibody-antigen complex structures, or their interacting residues, is still a laborious process. Several computational methods for predicting complex models or interface residues on antibody (paratope) or antigen (epitope) have been developed, but the problem of integrating these methods to archive a robust and coherent solution remains challenging. With the recent breakthroughs in protein structural modeling by Deep Learning [1-3], we revisit this important problem and assess the impact of state-of-the-art protein modeling on antibody-antigen docking and binding site prediction.

AbAdapt is a pipeline that combines antibody and antigen modeling with rigid docking and re-scoring in order to derive antibody-antigen specific features for epitope prediction [4]. As has been reported by others, the rigid docking and scoring steps are sensitive to the quality of the input models [5, 6]. By default, AbAdapt accepts sequences as input and uses Repertoire Builder, a high-throughput template-based method, for antibody modeling. However, AbAdapt can also accept structures as input for antibodies, antigens, or both. Here, we assessed the effect of using AlphaFold2 antibody models in the AbAdapt pipeline in a large-scale benchmark using leave-one-out cross validation (LOOCV) and also a large and diverse holdout set. In addition, the improved AbAdapt-AlphaFold2 (AbAdapt-AF) pipeline was tested using a set of recently determined antibodies that target various epitopes on a common antigen: the SARS-Cov-2 spike receptor binding domain (RBD). We found that the use of AlphaFold2 significantly improved the performance of AbAdapt, both at the level of protein structure and predicted binding sites.

## 2. Materials and methods

### 2.1 Datasets

The preparation of data for LOOCV and holdout tests was described previously [4]. In brief, crystal structures of antibody-antigen complexes were gathered from the Protein Data Bank (downloaded May. 24, 2021) [7]. We filtered the complexes using the following criteria: Complete heavy (H) and light (L) variable chain, antigen length equal to or greater than 60 amino acids and resolution less than 4 Å. Also, antigens that were themselves antibodies were removed. Antibody Complementarity-determining regions (CDR) sequences were annotated with the AHo numbering scheme [8] by the ANARCI program [9]. Next, the sequence redundancy was removed as follows: A pseudo-sequence of each antibody composed of heavy and light variable domain was constructed and clustered using CD-HIT [10] at an 85% identity threshold. Any sequences that resulted in failure by third-party software in subsequent steps were also removed. This filtering resulted in 720 non-redundant antibody-antigen model pairs. For training, LOOCV was performed using 620 randomly chosen antibody-antigen queries. The remaining 100 queries functioned as an independent holdout set for testing.

We also collected anti-SARS-Cov-2 RBD antibodies on Sept. 5, 2021 from the PDB [7]. Antibody pseudo-sequences were constructed as described above and clustered with the LOOCV training dataset using CD-HIT at an 85% sequence identity threshold. After superimposing all common epitopes for all pairs of antibodies, the C-alpha root-mean-square deviation (RMSD) was used to cluster the antibodies by single-linkage hierarchical clustering with a 10.0 Å cutoff. Within each cluster, the PDB entry with the most recent release date was chosen as the representative. This procedure resulted in the identification of 25 novel anti-RBD antibody-antigen complexes.

### 2.2 Epitope and Paratope definition

The epitope (paratope) residues were defined as any residue with at least one heavy atom within 5 Å of the antigen (antibody), as measured by Prody 2.0 [11].

### 2.3 Antibody modeling by AlphaFold2

The antibodies from the LOOCV and holdout sets were modeled independently from antigens using the full AlphaFold2 pipeline with default parameters [1]. We concatenated the sequences of H and L chains in the Fab region via a poly-glycine linker (32G). After cleaving the linker, the rank-one model was used in all subsequent calculations. Antibody CDRs were annotated as described in section 2.1. To evaluate the modeling accuracy of the CDRs, we first superimposed the +/- 4 flanking amino acid residues of each CDR in predicted models onto native antibody structures. Next, we calculated the RMSD of the CDRs by Prody 2.0 [11]. The paratope RMSD was computed similarly after superimposing the paratope of the model onto the native structure.

### 2.4 AbAdapt and AbAdapt-AF pipelines

AbAdapt was run using default parameters, as described previously [4]. In brief, antibodies were modeled using Repertoire Builder [12] and antigens were modeled using Spanner [13]. Here, any templates with bound antibodies overlapping with the true epitope were excluded. Two docking engines (Hex [14] and Piper [15]) were used to sample rigid docking degrees of freedom. Machine learning models were used to predict initial epitope and paratopes, score Hex poses, score Piper poses, score clusters of Hex and Piper poses, and predict antibody-specific epitope residues. For LOOCV calculations, each time an antibody-antigen pair was used as a query, each ML model was re-trained. For the holdout datasets, training was performed on the entire LOOCV dataset. For the AbAdapt-AF pipeline, the only procedural difference was that AlphaFold2 was used for antibody prediction instead of the default Repertoire Builder method. Naturally, all ML models were re-trained specifically for this use case.

### 2.5 RBD-antibody complex prediction

A model of SARS-Cov-2 RBD (329–532 aa) was built using Spanner and used as the antigen in all subsequent docking steps. The modeling of 25 anti-RBD antibodies was performed using AlphaFold2 on the ColabFold platform (v1.2) using the AlphaFold2-ptm model type with templates and Amber relaxation [16]. The rank-one antibody model was used in the downstream workflow of docking and complex modeling.

#### 2.5.1 ZDOCK workflow

The global protein-protein docking program ZDOCK (https://zdock.umassmed.edu/, v3.0.2) [17] was utilized with default parameters for antibody-RBD binding prediction. The top 10 models were retained for analysis.

#### 2.5.2 HawkDock workflow

HawkDock integrates the ATTRACT docking algorithm, the HawkRank scoring function, and the Molecular Mechanics/Generalized Born Surface Area free energy decomposition for protein-protein interface analysis [18]. We used the HawkDock server with default parameters (http://cadd.zju.edu.cn/hawkdock/) for antibody-RBD binding prediction. The top 100 models were retained for analysis.

#### 2.5.3 HDOCK workflow

An integrated template-based and template-free protein-protein docking program HDOCK (http://hdock.phys.hust.edu.cn/) [19] was utilized with template-free docking for antibody-RBD binding prediction. The top 100 models were retained for analysis.

#### 2.5.4 RBD-antibody complex evaluation

To evaluate the accuracy of predicted complexes, DockQ (https://github.com/bjornwallner/DockQ)

[20] was utilized with CAPRI criteria [21].

### 2.6 RBD epitope prediction

We used the following third-party methods for epitope prediction: DiscoTope2 (https://services.healthtech.dtu.dk/service.php?DiscoTope-2.0), which predicts discontinuous B-cell epitopes [22]; BepiPred2 (https://services.healthtech.dtu.dk/service.php?BepiPred-2.0), which uses a random forest regression algorithm [23]; EpiDope (https://github.com/flomock/EpiDope), which uses a deep neural network to predict linear B-cell epitopes based on antigen amino acid sequence [24]. For binary labeling of the predicted epitopes, the default thresholds of 0.5 for BepiPred2 [23] and 0.818 for EpiDope [24] were used. To predict the structural epitopes specific to a given antibody, EpiPred (http://opig.stats.ox.ac.uk/webapps/newsabdab/sabpred/epipred/) was used with 25 anti-RBD antibody models from AlphaFold2 and an RBD model from Spanner [25]. The rank-one epitope prediction from EpiPred was chosen for all subsequent analysis. For AbAdapt-AF and AbAdapt, a threshold of 0.5 for final epitope prediction was used [4]. The performance indices of epitope prediction were calculated using the Python package Scikit-learn (v 0.23.2) [26].

### 2.7 Statistical analysis and Visualization

Wilcoxon matched-pairs signed rank test and descriptive analysis were performed using GraphPad Prism 8 (GraphPad Software, San Diego, CA). For graphing, the Python package Seaborn (v 0.11.0) and Matplotlib (v 3.3.1) were used. We visualized the RBD-antibody complexes and RBD structure with its corresponding epitope map using PyMOL (The PyMOL Molecular Graphics System, v 2.3.3 Schrödinger, LLC.)

## 3. Results

### 3.1 Improvement in antibody modeling using AlphaFold2

The CDRs constitute the greatest source of sequence and structural variability in antibodies and also largely overlap with their paratope residues. Here, we systematically evaluated the performance of antibody variable region structural models by Repertoire Builder and AlphaFold2 using the LOOCV and holdout datasets. The accuracy of antibody modeling improved significantly in both the LOOCV (Supplementary Fig. 1A) and holdout (Supplementary Fig. S1B) sets. The improvement of AlphaFold2 over Repertoire Builder was particularly apparent in the modeling of the most challenging CDR loop, CDR-H3: the average RMSD by AlphaFold2 for the LOOCV set dropped from 4.38 Å to 3.44 Å, a 21.50% improvement over Repertoire Builder (Supplementary Table. 1). Similar results were obtained for the holdout set (4.44 Å to 3.62 Å, a 18.43% improvement).

As expected, the improvement of CDR loop modeling by AlphaFold2 resulted in improved paratope modeling: the paratope RMSD dropped from 2.69 Å to 2.08 Å (a 22.73% improvement) in the LOOCV set and from 2.83 Å to 2.12 Å (a 25.26% improvement) in the holdout set (Fig. 1A and Supplementary Table. 1). Using a threshold of paratope RMSD > 4 Å to define low-quality models, the ratio of low-quality models by AlphaFold2 dropped from 14.35% to 7.74% in the LOOCV set and 21.0% to 11.0% in the holdout set, approximately a 2-fold decrease. These improvements are of interest in antibody-antigen docking because we previously observed that AbAdapt docking performance was sensitive to paratope structural modeling errors [4].

**Figure 1.**
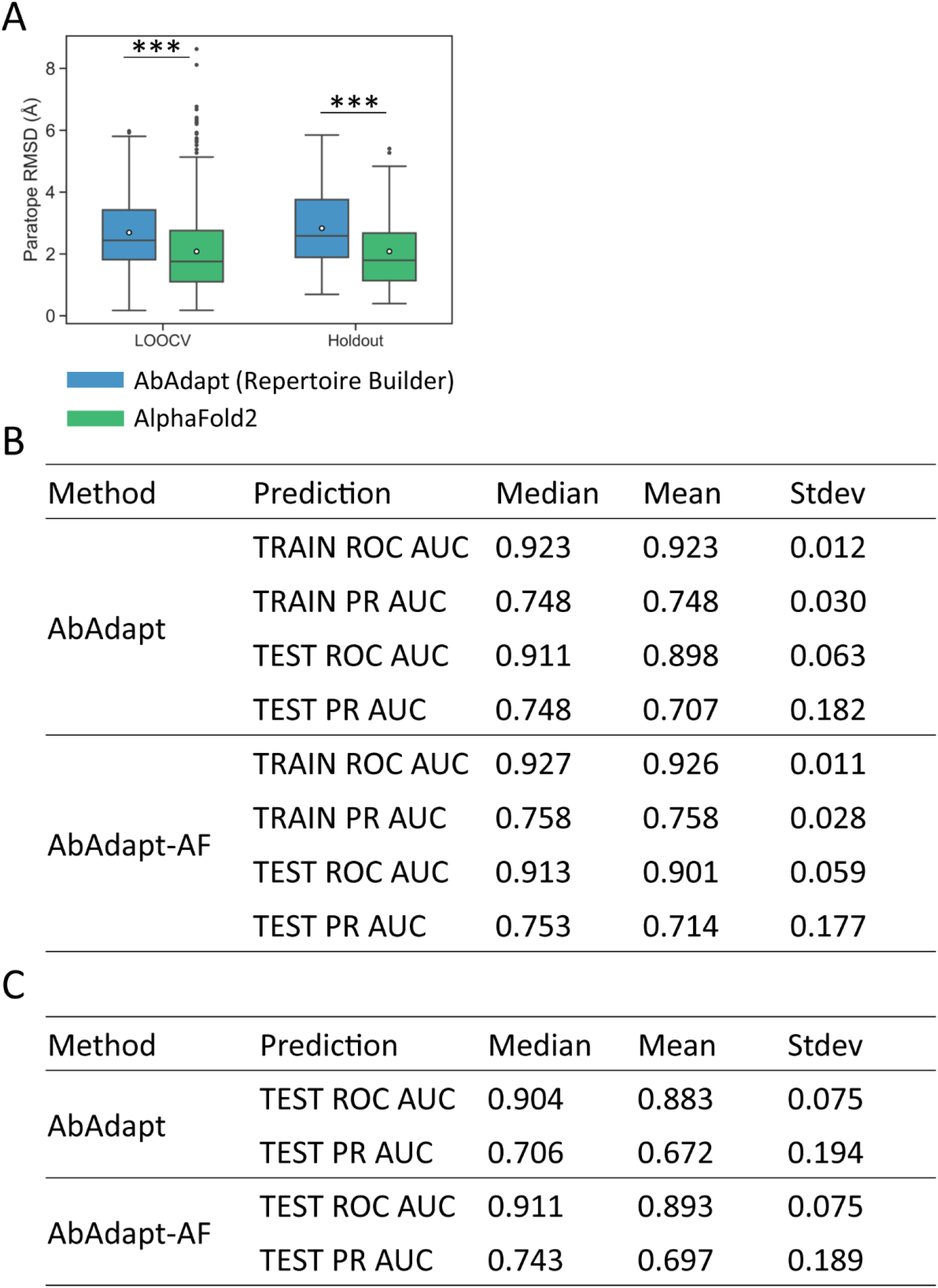
Comparison of the performance of docking and paratope prediction between AbAdapt and AbAdapt-AF. (A) The paratope RMSD of antibody model in LOOCV training set and Holdout set by AbAdapt or AlphaFold2. Wilcoxon matched-pairs signed rank test was performed to compare the corresponding performance between AbAdapt and AbAdapt-AF (\*\*\**P*≤ 0.001). The empty circle in each box indicated the average value. Comparison of the paratope prediction performance of antibodies by AbAdapt and AlpahFold2 in LOOCV set (B) and holdout set (C).

### 3.2 Improvement of AbAdapt pipeline using AlphaFold2 antibody models

After including the more accurate AlphaFold2 antibody models in the AbAdapt pipeline, we evaluated the influence on each of the main steps: paratope prediction, initial epitope prediction, docking, scoring Piper-Hex clusters and antibody-specific epitope prediction.

#### 3.2.1 Paratope prediction

The Area Under the Curve (AUC) of Receiver Operating Characteristic curve (ROC) and Precision-Recall (PR) curves for paratope prediction for the LOOCV dataset were calculated for AbAdapt and AbAdapt-AF (Fig. 1B). We found close median ROC AUCs for both training (0.927) and testing (0.913) by AbAdapt-AF which showed slight improvement over AbAdapt (0.923 for training and 0.911 for testing). The median PR AUC for testing improved from 0.748 (AbAdapt) to 0.753 (AbAdapt-AF). In the holdout set, the median ROC AUC for paratope prediction by AbAdapt-AF was 0.911, which was close to that of the LOOCV set (0.913) (Fig. 1B and 1C). A dramatic improvement of median PR AUC was also observed: from 0.706 (AbAdapt) to 0.743 (AbAdapt-AF) (Fig .1C). Thus, introducing more accurate antibody models from AlphaFold2 clearly improved paratope prediction.

#### 3.2.2 Initial epitope prediction

In the case of initial epitope prediction, AbAdapt-AF exhibited a median test ROC AUC of 0.694 in the LOOCV set and 0.695 in the holdout set which was identical to AbAdapt, as expected, since both pipelines used the same antigen model from Spanner for training. Meanwhile, these values were far lower than the median training ROC AUC: 0.863 (Supplementary Table. 2). The difference between training and testing suggests that the initial epitope predictor does not generalize well to unseen data and highlights the inherent limitation in epitope prediction without reference to a specific antibody. This issue is addressed in section 3.2.5 by performing epitope prediction with specific antibody-derived features.

#### 3.2.3 Hex and Piper docking

We first analyzed the effect of AlphaFold2 models on the frequency of “true” poses produced by Hex or Piper in the LOOCV set. Here, as in our previous work [4], a relatively loose cutoff value of 15 Å for the RMSD of the interface residues (IRMSD), along with epitope and paratope accuracies of 50%, was used to define a “true” pose. The median true pose ratio of the Hex engine increased modestly from 1.50% to 1.53% (Supplementary Fig. 2C), which was not statistically significant (Supplementary Fig. 2A). This is somewhat expected, since Hex was used for local docking, guided by the initial epitope predictions, which themselves were not significantly affected by the use of AlphaFold2 antibody models. In contrast to Hex, the median Piper true pose ratio, which increased from 1.43% to 1.57%, significantly improved upon use of AlphaFold2 antibody models (Supplementary Fig. 2A and 2C). For the holdout set, the median true pose ratio for both Hex and Piper were not significantly improved by use of AlphaFold2 (Supplementary Fig. 3A and 3C). These results indicate that simply supplying better models does not guarantee improved docking. A similar observation was reported in a recent study using AlphaFold2 models in the ClusPro web server (Supplementary Fig. 4) [27].

#### 3.2.4 Combined Piper-Hex clustering and scoring

The top Piper and Hex poses from the docking steps were co-clustered and re-scored. Here, we observed an improvement in the AbAdapt-AF pipeline compared with AbAdapt. In the LOOCV set, the median clustered true pose ratio of Hex improved from 1.16% to 1.31% while that for Piper improved 2.32% to 2.78%, resulting in a significantly improved median combined Hex-Piper true pose ratio (2.42% versus 2.83%) for AbAdapt versus AbAdapt-AF, respectively (Supplementary Fig. 2B and 2C). The median rank of the best true pose (with 1 being perfect) dropped significantly from 17 to 13.5, a 20.59% improvement (Supplementary Fig. 2D). We next imposed a stricter (7 Å) IRMSD cutoff; even under this stricter criterion, the median true rank dropped from 54.5 to 43, a 21.10% improvement (Supplementary Fig. 2D). This improvement was also reflected in the ratio of queries with true poses under the stricter 7 Å IRMSD cutoff (success ratio) in the LOOCV set: The AbAdapt-AlphaFold2 success ratio was 55.81%, compared with 46.67% for AbAdapt (Fig. 2A).

**Figure 2.**
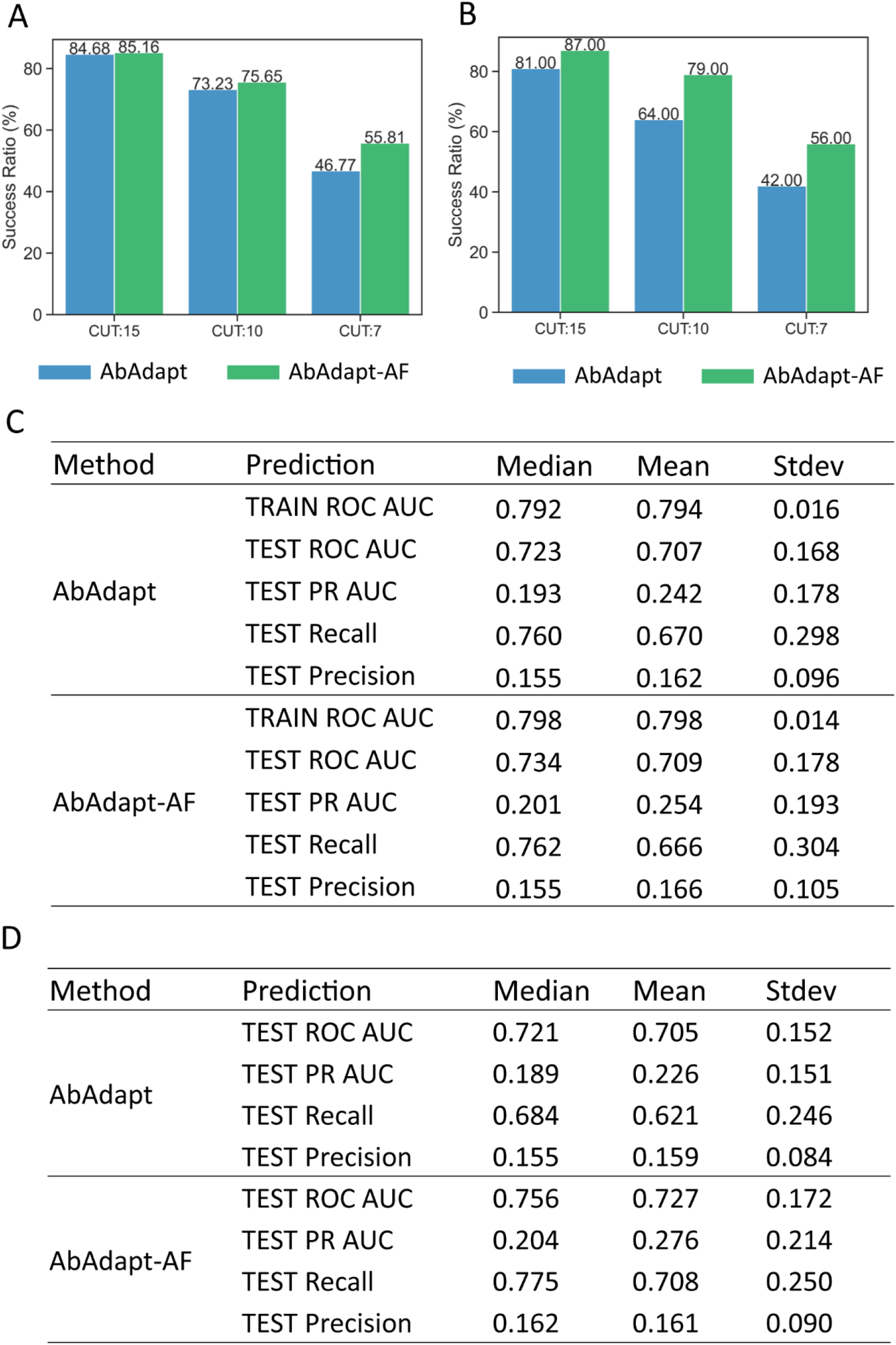
Comparison of the performance of pose clustering and epitope prediction between AbAdapt and AbAdapt-AF. The success ratio of pose clustering in LOOCV set (A) and holdout set (B) after clustering and combining the pose from Piper and Hex. When setting the true cutoff of interface RMSD (RCUT) 15, 10 and 7 Å, the corresponding success ratio was shown above each bar. The final epitope prediction performance in LOOCV set (C) and holdout set (D) after introducing the post-docking features of specific antibodies.

For the unseen holdout set, the median clustered true pose ratio for Hex improved but was not statistically significant: 0.56 +/- 8.15 % (AbAdapt) versus 1.25 +/- 6.69% (AbAdapt-AF) (Supplementary Fig. 3B and 3C). However, the median clustered true pose ratio of Piper improved significantly: 2.18 +/- 3.97% (AbAdapt) versus 2.66 +/- 3.72% (AbAdapt-AF), resulting in a significantly improved median combined Hex-Piper true pose ratio (2.18 +/- 4.12% versus 2.7 +/- 3.9%) for AbAdapt versus AbAdapt- AF. AbAdapt-AF outperformed AbAdapt in all three IRMSD cutoff values and the success ratio was very close to that of the LOOCV set (Fig. 2A and 2B). Taken together, the performance of scoring and clustering of Hex and Piper poses significantly improved by using AbAdapt-AF.

#### 3.2.5 Antibody-specific epitope predictions

After retraining the epitope predictor with docking features, we observed significant improvement in antibody-specific epitope prediction. In the LOOCV set, the median test ROC AUC increased from 0.694 (initial epitope prediction) to 0.734 (antibody-specific epitope prediction) by AbAdapt-AF, compared with 0.694 (initial epitope prediction) to 0.723 (antibody-specific epitope prediction) by AbAdapt (Fig. 2C and Supplementary Table. 2). An even greater improvement was observed in the holdout set: AbAdapt-AF, the median test ROC AUC improved from 0.695 (initial epitope prediction) to 0.756 (antibody-specific epitope prediction); for AbAdapt, the corresponding values were 0.695 (initial epitope prediction) and 0.721 (antibody-specific epitope prediction), as shown in Fig. 2D and Supplementary Table. 2. There are relatively few antigen residues that make up the epitope (class imbalance of epitope and non-epitope residues). The PR ROC baseline is given by the ratio Ntrue/Nfalse (number of epitope residues/number of non-epitope residues). By this definition, the median PR AUC baseline was 0.094 in the holdout set (Supplementary Fig. 5). In the holdout set, the median PR AUC of the antibody-specific epitope prediction reached 0.204 (a 117.02% improvement over the baseline) by AbAdapt-AF, while the corresponding value was 0.189 for AbAdapt, representing a smaller (101.06%) improvement over the baseline (Fig. 2D). This result supports the conclusion that the inclusion of AlphaFold2 significantly improved the final epitope predictions, even for inputs that have never been seen by the ML models.

We note that, in the current AbAdapt-AF pipeline, antibody models were built by AlphaFold2 while antigen models were built by the template-based modeling engine Spanner. To investigate the sensitivity of performance to the antigen models, we repeated the antigen modeling using AlphaFold2. Here antibody-specific epitope prediction median test ROC AUC values were 0.745 (0.721 by AbAdapt) and median test PR AUC values were 0.195 (0.189 by AbAdapt) (Supplementary Table. 3). Compared to the median test ROC AUC of the antibody-specific epitope prediction using AbAdapt (0.721), AbAdapt-AF achieved better performance, using either Spanner antigen models (0.756) or AlphaFold2 antigen models (0.745) as input. These results suggest that the main impact on AbAdapt was due to the use of AlphaFold2 for antibody modeling.

### 3.3 Anti-RBD docking performance

To assess AbAdapt on a realistic test case, we prepared 25 non-redundant SARS-Cov-2 anti-RBD antibodies that were not among the training data used by AbAdapt or AlphaFold. These antibodies targeted 16 epitope clusters in the RBD (Supplementary Fig. 6). We next assessed the performance of AbAdapt-AF along with ZDOCK, AbAdapt, HawkDock, and HDOCK, as shown in Fig. 3A and 3B. In terms of rank1 models, AbAdapt-AF, a total of 4 (16%) acceptable or better models were produced by AbAdapt-AF. The corresponding success rates for the other methods were: ZDOCK (1), AbAdapt (0), HawkDock (0), HDOCK (1). Similarly, in terms of 100 top-ranked models, a total of 15 (60%) acceptable or better models were built by AbAdapt-AF. The corresponding success rates of the other tested methods were: AbAdapt (8), HawkDock (8), HDOCK (10). AbAdapt-AF performed better than AbAdapt to a similar degree that described in the LOOCV and holdout assessment. Considering all cluster representatives (mean value 539 per query), by CAPRI criteria (IRMSD < 10 Å), the AbAdapt-AF success ratio was 84% (21 queries) which was close to 75.65% in the LOOCV and 79% in the holdout set (Fig. 2A, 2B and 3B).

**Figure 3.**
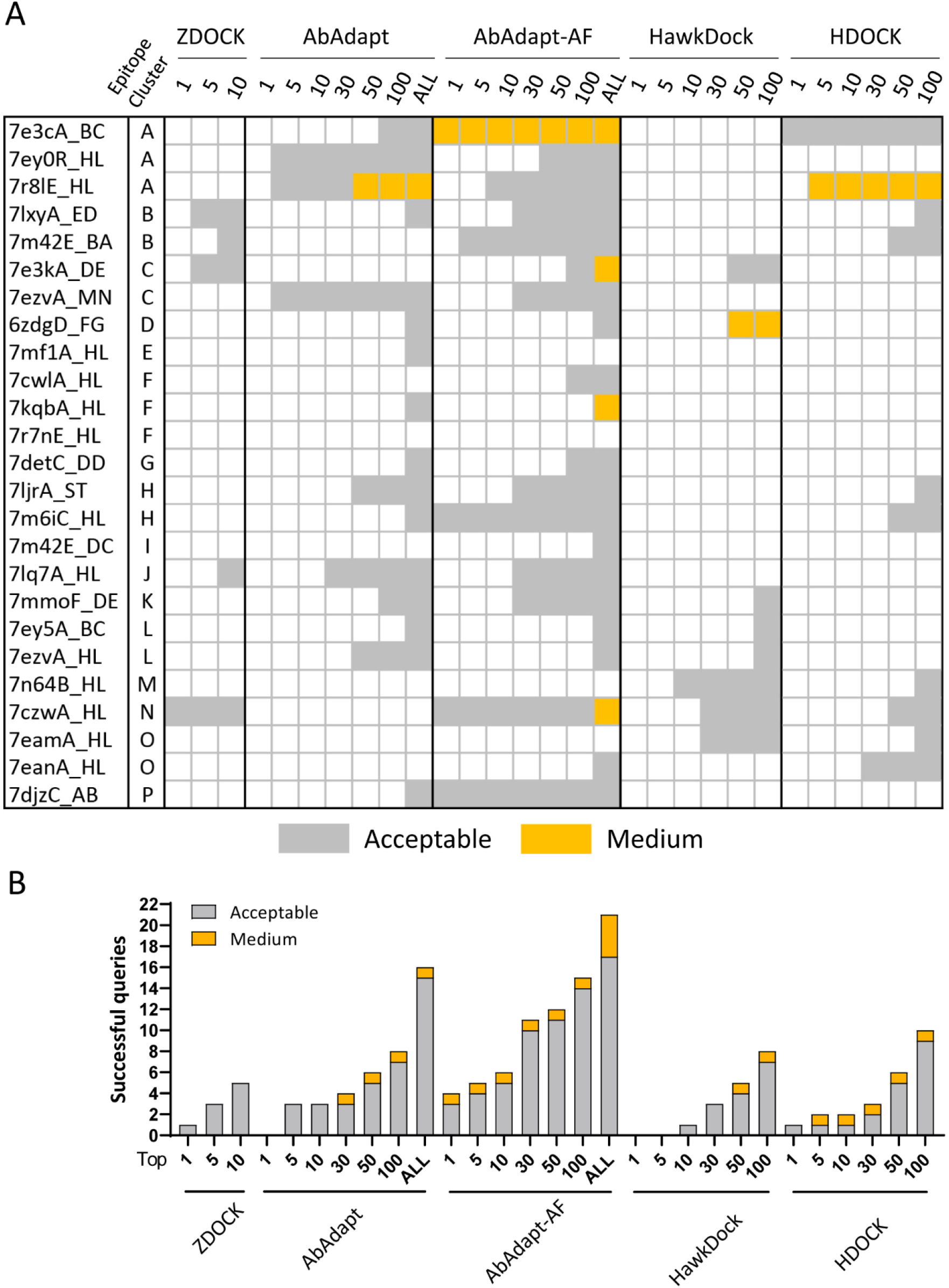
Docking performance of 25 anti-RBD antibody complexes. (A) The CAPRI quality of the best model from top 1/5/10 ranked models by ZDOCK, top 1/5/10/30/50/100/all ranked models by AbAdapt and AbAdapt-AF, and top 1/5/10/30/50/100 by HawkDock and HDOCK. The color of each cell was their corresponding CAPRI quality showed as acceptable (grey) and medium (orange). (B) The number of successful queries by ZDOCK, AbAdapt, AbAdapt-AF, HawkDocK, HDOCK.

### 3.4 Antibody-specific RBD epitope predictions

We next analyzed epitope prediction in the RBD benchmark. In addition to using Abadapt and Abadapt-AF, we also evaluated epitope prediction tools that are not based on docking. These included: BepiPred2 for linear epitope prediction, DiscoTope2 for structural epitope prediction, EpiPred for antibody-specific epitope prediction, and EpiDope, which uses a deep neural network based on antigen sequence features. AbAdapt-AF achieved the highest ROC AUC (0.793) and PR AUC (0.411) using the antibody-specific epitope prediction probability (Fig. 4A). We found the average AbAdapt-AF ROC AUC (0.793) of the RBD benchmark was close to the values in LOOCV (0.709) and holdout (0.727) runs (Fig. 2C and 2D). The improvement of average ROC AUC in the RBD benchmark (7.02%) was similar to that observed in holdout set (4.85%) by AbAdapt-AF (Fig. 2D and 4A), thus again demonstrating the robustness of antibody-specific epitope prediction using AbAdapt-AF. Meanwhile, AbAdapt-AF achieved the highest precision (0.324), F1 score (0.378) and MCC (0.325) among the six tested methods when setting a fixed epitope probability threshold (Fig. 4B) as described in section 2.6.

**Figure 4.**
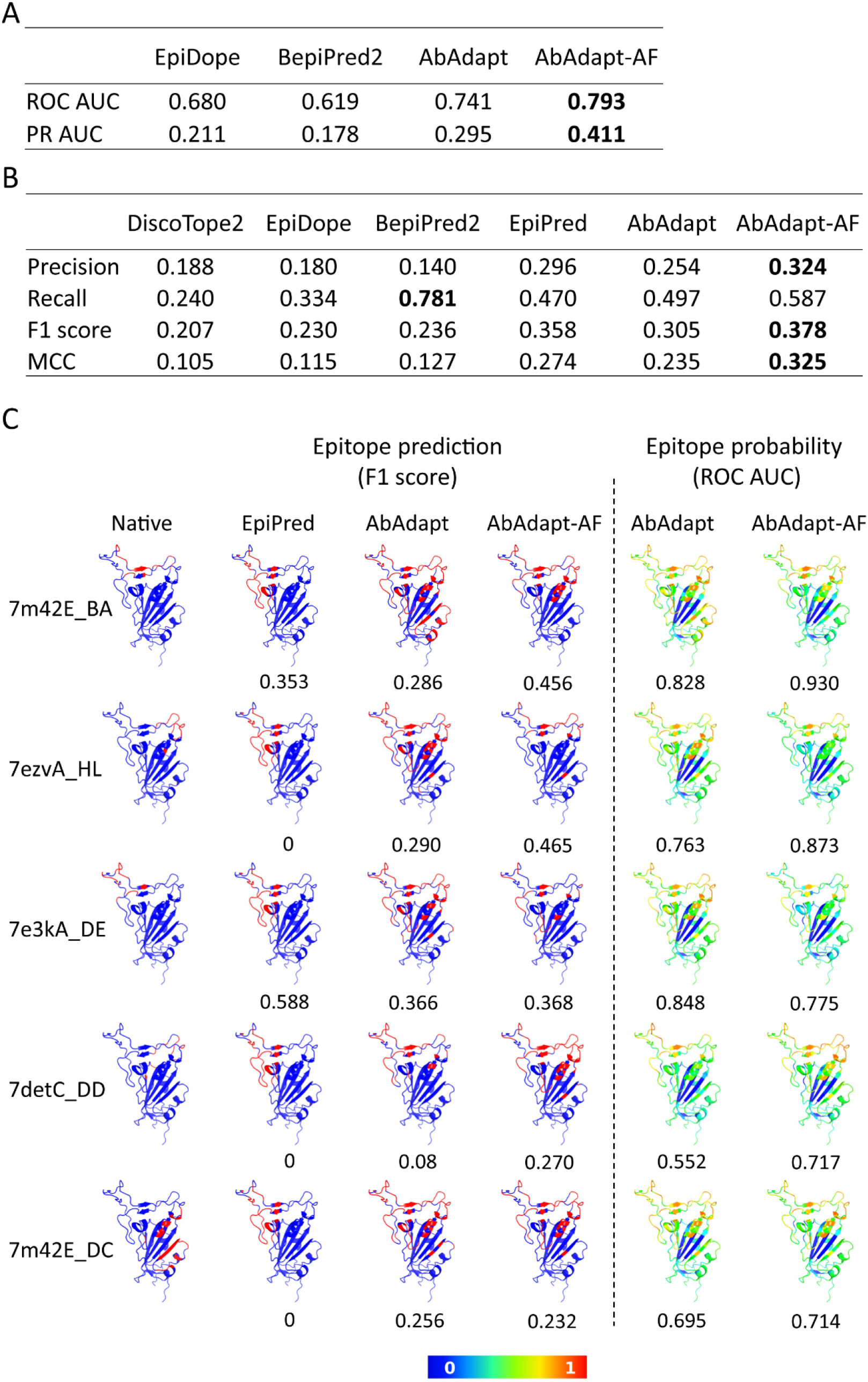
Epitope prediction of 25 anti-RBD antibody complexes. The comparison of epitope prediction performance using probability (A) and a threshold for epitope classification (B). The performance indices are calculated in evaluation metrics and averaged. Bold character indicated the highest value of each item. (C) Epitope map visualization of representative queries. The native epitope (column 1) and predicted epitope by EpiPred (column 2), AbAdapt (column 3), and AbAdapt-AF (column 4) are colored red on the RBD surface. The probability of prediction by AbAdapt and AbAdapt-AF are shown in columns 5 and 6. The value below each prediction is indicated as F1 score (left) and ROC AUC (right).

The prediction performances of DiscoTope2, EpiDope, and BepiPred2, which do not take antibody into consideration, were systematically lower than those of the other methods. We noticed that BepiPred2 achieved the highest recall (0.781), but this value was obtained at a precision of only 0.140. Although EpiPred considers antibody information, the results were not very sensitive to the antibodies in this set, as all epitope predictions were in the same region of the RBD (Fig. 4C).

## 4. Discussion

In this study, we demonstrated that introducing a more accurate antibody model in the AbAdapt antibody-specific epitope prediction pipeline had a significantly positive effect at various levels. Nevertheless, there is still much room for improvement. Considering the RBD benchmark, we evaluated six methods for epitope prediction on the SARS-CoV-2 spike RBD, an antigen that can be targeted by one or more antibodies on almost its entire molecular surface [28]. We found AbAdapt-AF showed higher epitope prediction accuracy than tested methods in this benchmark. Nevertheless, the ROC AUC (0.793) and precision (0.324) values imply that many false-positive epitope predictions are expected. To compare the epitope prediction performance of AbAdapt-AF and other tools fairly and systematically, a broader benchmark containing various antigens will be needed. At present, the use of AlphaFold2 in the context of AbAdapt suggests that this combination represents a robust but incremental way of improving antibody-specific epitope predictions. Newly released antibody modeling tools such as ABlooper [2] and DeepAb [3] showed better performance than AlphaFold2 in the Rosetta Antibody Benchmark [2], and these methods should also be considered when running AbAdapt with structural models as input.

## Acknowledgments

We would like to thank all members of the Systems Immunology Lab for very helpful discussion. The computation was performed using Research Center for Computational Science, Okazaki, Japan (Project: 21-IMS-C116 and 22-IMS-C136).

## Highlights

AlphaFold2 can produce a more accurate antibody models than Repertoire Builder, which is used by default in the AbAdapt pipeline.

Integration of AlphaFold2 in AbAdapt (AbAdapt-AF) resulted in improved paratope prediction, clustering, re-ranking of pose clusters, and antibody-specific epitope predictions compared with the default AbAdapt pipeline.

AbAdapt-AF outperformed other three tested docking methods, resulting in an 60% success ratio (queries with at least one correctly predicted pose in the top 100) in the anti-RBD antibody complex benchmark.

AbAdapt-AF demonstrated higher epitope prediction accuracy than other tested tools in the anti-RBD antibody complex benchmark.

