## Supplemental Materials for "Improved antibody-specific epitope prediction using AlphaFold and AbAdapt"

**This file includes:**

Supplementary Table 1. The RMSD (Å) of CDRs and paratope in the LOOCV and holdout set.

Supplementary Table 2. Comparison of initial epitope prediction between AbAdapt and AbAdapt-AF in LOOCV and holdout set.

Supplementary Table 3. Comparison of the performance of antibody-specific epitope prediction between AbAdapt and AbAdapt-AF.

Supplementary Figure 1. Comparison of the modeling performance of antibodies between AbAdapt and AlpahFold2.

Supplementary Figure 2. Comparison of docking performance and pose combination between AbAdapt and AbAdapt-AF in LOOCV set.

Supplementary Figure 3. Comparison of docking performance and pose combination between AbAdapt and AbAdapt-AF in holdout set.

Supplementary Figure 4. The performance of antigen-antibody complex prediction by AlphaFold2.

Supplementary Figure 5. PR ROC baseline of LOOCV and holdout sets.

Supplementary Figure 6. The visualization of 25 SARS-Cov-2 RBD-antibody complexes.

The RMSD of CDRs and paratope in the LOOCV set

| | | CDR-H1 | CDR-H2 | CDR-H3 | CDR-L1 | CDR-L2 | CDR-L3 | Paratope |
|---|---|---|---|---|---|---|---|---|
| AlphaFold2 | Median | 0.61 | 0.64 | 2.88 | 0.59 | 0.49 | 0.78 | 1.76 |
| | Mean | 0.93 | 0.89 | 3.44 | 0.84 | 0.64 | 1.15 | 2.08 |
| | Stdev | 0.89 | 0.98 | 2.42 | 0.90 | 0.82 | 1.08 | 1.28 |
| Repertoire Builder | Median | 0.81 | 0.81 | 4.01 | 0.67 | 0.57 | 0.92 | 2.44 |
| | Mean | 1.14 | 1.07 | 4.38 | 0.95 | 0.72 | 1.37 | 2.69 |
| | Stdev | 0.97 | 1.03 | 2.31 | 0.90 | 0.79 | 1.24 | 1.20 |

The RMSD of CDRs and paratope in the holdout set

| | | CDR-H1 | CDR-H2 | CDR-H3 | CDR-L1 | CDR-L2 | CDR-L3 | Paratope |
|---|---|---|---|---|---|---|---|---|
| AlphaFold2 | Median | 0.67 | 0.74 | 2.86 | 0.62 | 0.55 | 0.25 | 1.80 |
| | Mean | 1.14 | 1.04 | 3.62 | 0.88 | 0.60 | 1.31 | 2.12 |
| | Stdev | 1.06 | 0.86 | 2.88 | 1.01 | 0.41 | 1.17 | 1.17 |
| Repertoire Builder | Median | 0.85 | 1.07 | 4.07 | 0.72 | 0.61 | 0.28 | 2.59 |
| | Mean | 1.34 | 1.31 | 4.44 | 1.05 | 0.74 | 1.53 | 2.83 |
| | Stdev | 1.12 | 1.01 | 2.19 | 1.09 | 0.79 | 1.15 | 1.18 |

**Supplementary Table 1. The RMSD (Å) of CDRs and paratope in the LOOCV and holdout set.**

Initial epitope prediction of LOOCV set

| | Prediction | Median | Mean | Stdev |
|---|---|---|---|---|
| AbAdapt | TRAIN ROC AUC | 0.863 | 0.854 | 0.048 |
| | TEST ROC AUC | 0.694 | 0.687 | 0.147 |
| | TEST PR AUC | 0.165 | 0.200 | 0.137 |
| | TEST Recall | 0.625 | 0.599 | 0.241 |
| | TEST Precision | 0.149 | 0.157 | 0.088 |
| AbAdapt-AF | TRAIN ROC AUC | 0.863 | 0.854 | 0.048 |
| | TEST ROC AUC | 0.694 | 0.687 | 0.147 |
| | TEST PR AUC | 0.165 | 0.200 | 0.137 |
| | TEST Recall | 0.625 | 0.599 | 0.241 |
| | TEST Precision | 0.149 | 0.157 | 0.088 |

Initial epitope prediction of holdout set

| | Prediction | Median | Mean | Stdev |
|---|---|---|---|---|
| AbAdapt | TEST ROC AUC | 0.695 | 0.690 | 0.146 |
| | TEST PR AUC | 0.162 | 0.212 | 0.159 |
| | TEST Recall | 0.571 | 0.593 | 0.224 |
| | TEST Precision | 0.150 | 0.159 | 0.092 |
| AbAdapt-AF | TEST ROC AUC | 0.695 | 0.690 | 0.146 |
| | TEST PR AUC | 0.162 | 0.212 | 0.159 |
| | TEST Recall | 0.571 | 0.593 | 0.224 |
| | TEST Precision | 0.150 | 0.159 | 0.092 |

**Supplementary Table 2. Comparison of initial epitope prediction between AbAdapt and AbAdapt-AF in LOOCV and holdout set.**

| | Prediction | Median | Mean | Stdev |
|---|---|---|---|---|
| AbAdapt | TEST ROC AUC | 0.721 | 0.705 | 0.152 |
| | TEST PR AUC | 0.189 | 0.226 | 0.151 |
| | TEST Sensitivity | 0.684 | 0.621 | 0.246 |
| | TEST Precision | 0.155 | 0.159 | 0.084 |
| AbAdapt-AF | TEST ROC AUC | 0.756 | 0.727 | 0.172 |
| | TEST PR AUC | 0.204 | 0.276 | 0.214 |
| | TEST Sensitivity | 0.775 | 0.708 | 0.250 |
| | TEST Precision | 0.162 | 0.161 | 0.090 |
| AbAdapt-AF (AF's Ab/Ag models) | TEST ROC AUC | 0.745 | 0.715 | 0.168 |
| | TEST PR AUC | 0.195 | 0.259 | 0.206 |
| | TEST Sensitivity | 0.771 | 0.691 | 0.258 |
| | TEST Precision | 0.158 | 0.160 | 0.096 |

**Supplementary Table 3. Comparison of the performance of antibody-specific epitope prediction between AbAdapt and AbAdapt-AF.** Analysis of the antibody-specific epitope prediction performance of the AbAdapt-AF pipeline that trained by antigen model from Spanner and antibody model from AlphaFold2 versus using antibody and antigen model both from AlphFold2 as input.

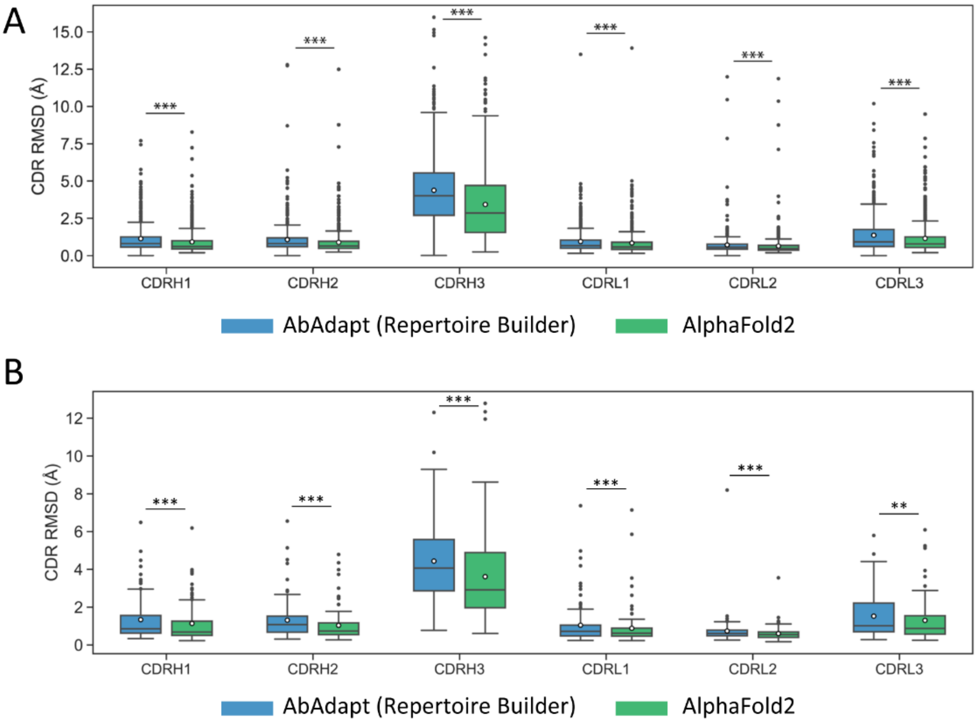


**Supplementary Figure 1. Comparison of the modeling performance of antibodies between AbAdapt and AlpahFold2.** The RMSD of six CDRs of antibody model in LOOCV training set with 620 queries (A) and Holdout set with 100 queries (B) by AbAdapt powered by Repertoire Builder (blue) or AlphaFold2 (green). The Wilcoxon matched-pairs signed rank test was performed to compare the corresponding performance between AbAdapt and AbAdapt-AF (***P*≤ 0.01; ****P*≤ 0.001). The empty circle in each box indicated the average value.

A

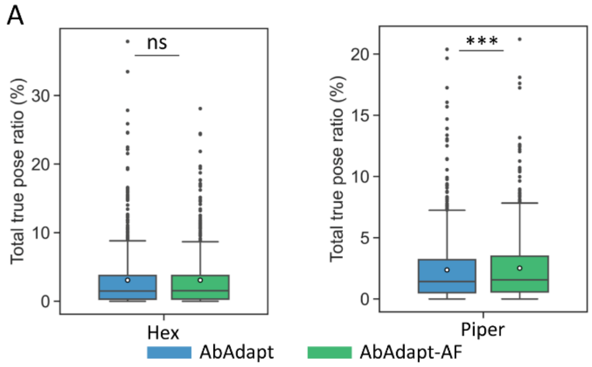


B

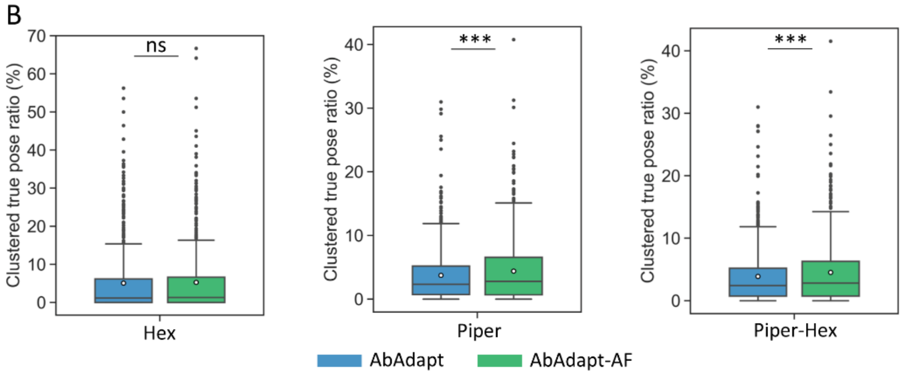


C

| | Hex true pose ratio (%) | | | Piper true pose ratio (%) | | | Hex clustered true pose ratio (%) | | | Piper clustered true pose ratio (%) | | | Combined Hex-Piper true pose ratio (%) | | |
|---|---|---|---|---|---|---|---|---|---|---|---|---|---|---|---|
| | Mean | Median | Stdev | Mean | Median | Stdev | Mean | Median | Stdev | Mean | Median | Stdev | Mean | Median | Stdev |
| AbAdapt | 3.09 | 1.5 | 4.63 | 2.37 | 1.43 | 2.85 | 5.08 | 1.16 | 9.03 | 3.73 | 2.32 | 5.05 | 3.88 | 2.42 | 5.24 |
| AbAdapt-AF | 3.08 | 1.53 | 4.23 | 2.51 | 1.57 | 2.88 | 5.27 | 1.31 | 8.68 | 4.4 | 2.78 | 4.39 | 4.52 | 2.83 | 4.55 |

D

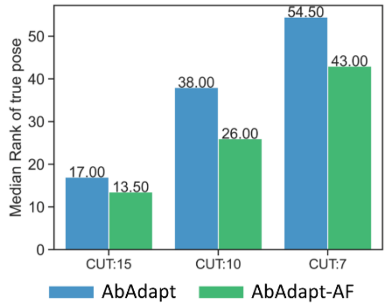


**Supplementary Figure 2. Comparison of docking performance and pose combination between AbAdapt and AbAdapt-AF in LOOCV set.** (A) The total true pose that produced by Hex (left) and Piper (right) by AbAdapt and AbAdapt-AF. (B) The true pose ratio after clustering the pose from Hex (left) and Piper (middle) separately and a combination of them (right). The Wilcoxon matched-pairs signed rank test was performed to compute the significance (****P*≤ 0.001). (C) The average and median values of each true pose ratio that related to (A) and (B). (D) The median rank of true poses after the combination of Hex-Piper clusters for the sharing successful queries among AbAdapt and AbAdapt-AF.

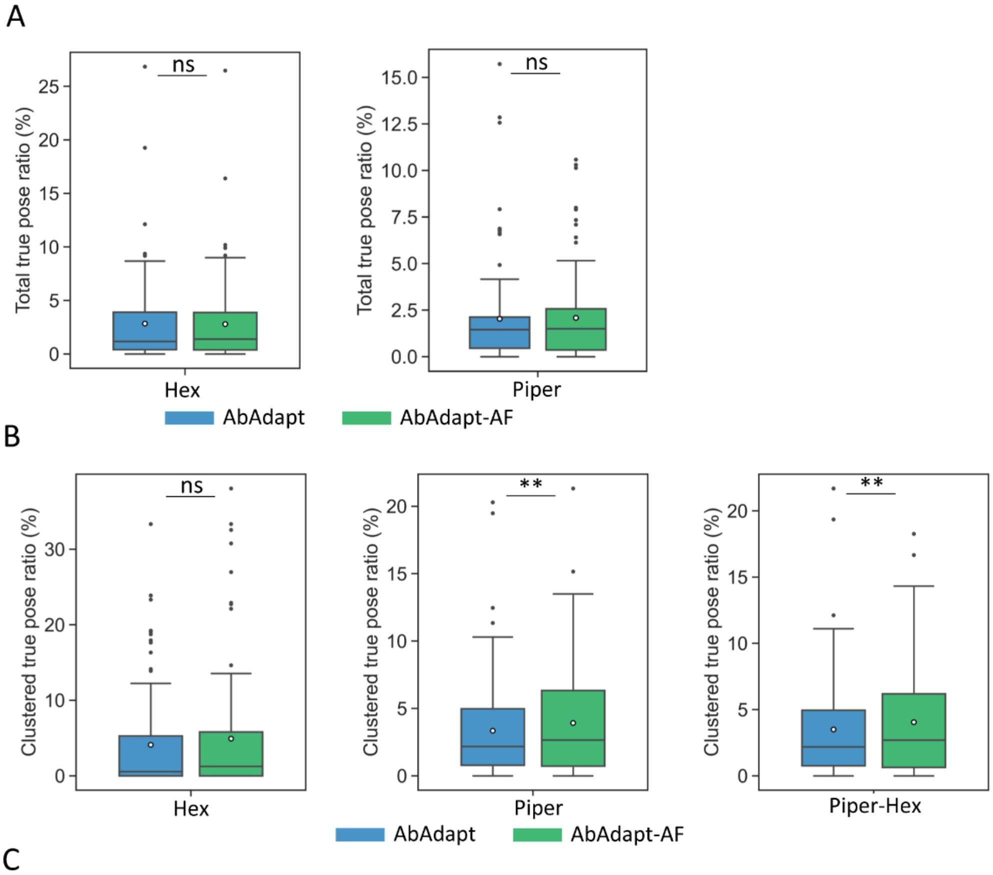


C

| | Hex true pose ratio (%) | | | Piper true pose ratio (%) | | | Hex clustered true pose ratio (%) | | | Piper clustered true pose ratio (%) | | | Combined Hex-Piper true pose ratio (%) | | |
|---|---|---|---|---|---|---|---|---|---|---|---|---|---|---|---|
| | Mean | Median | Stdev | Mean | Median | Stdev | Mean | Median | Stdev | Mean | Median | Stdev | Mean | Median | Stdev |
| AbAdapt | 2.83 | 1.17 | 3.82 | 2.03 | 1.45 | 2.34 | 4.11 | 0.56 | 8.15 | 3.35 | 2.18 | 3.97 | 3.5 | 2.18 | 4.12 |
| AbAdapt-AF | 2.79 | 1.39 | 4.02 | 2.09 | 1.5 | 2.7 | 4.92 | 1.25 | 6.69 | 3.92 | 2.66 | 3.72 | 4.05 | 2.7 | 3.9 |

**Supplementary Figure 3. Comparison of docking performance and pose combination between AbAdapt and AbAdapt-AF in holdout set.** (A) The total true pose that produced by Hex (left) and Piper (right) by AbAdapt and AbAdapt-AF. (B) The true pose ratio after clustering the pose from Hex (left) and Piper (middle) separately and a combination of them (right). The Wilcoxon matched-pairs signed rank test was performed to compute the significance (***P*≤ 0.01). (C) The average and median values of each true pose ratio that related to (A) and (B).

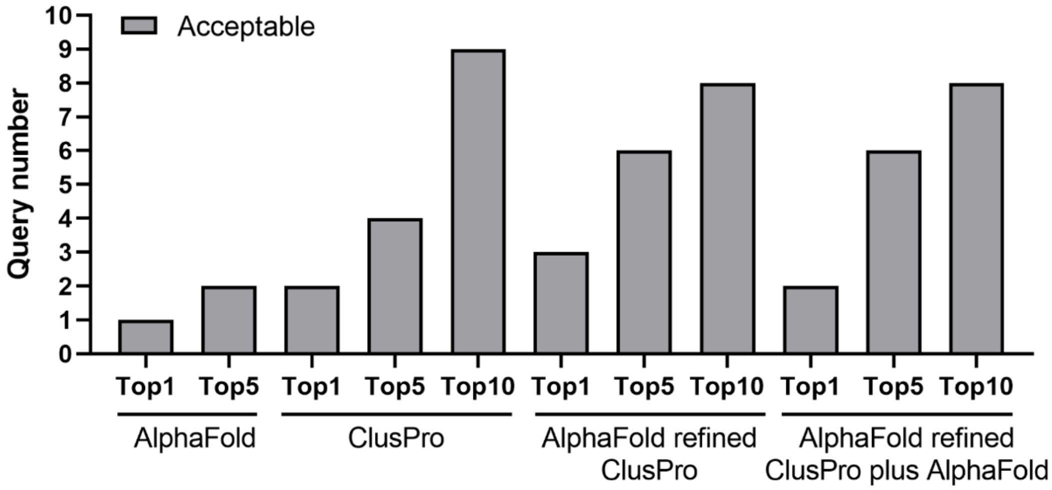


**Supplementary Figure 4. The performance of antigen-antibody complex prediction by AlphaFold2.** Reanalyzed the prediction accuracy of 32 Ag-Ab complexes by CAPRI criteria from a previous study (Ghani et al).

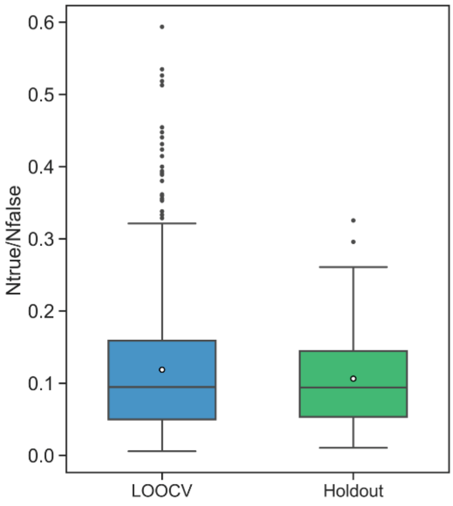


**Supplementary Figure 5. PR ROC baseline of LOOCV and holdout sets.** The baseline is based on the ratio of epitope and non-epitope amino acid residue. The “Ntrue” indicate the number of epitope residue and “Nfalse” indicate the number of non-epitope residue in each antigen from the LOOCV set (blue) and the holdout set (green).

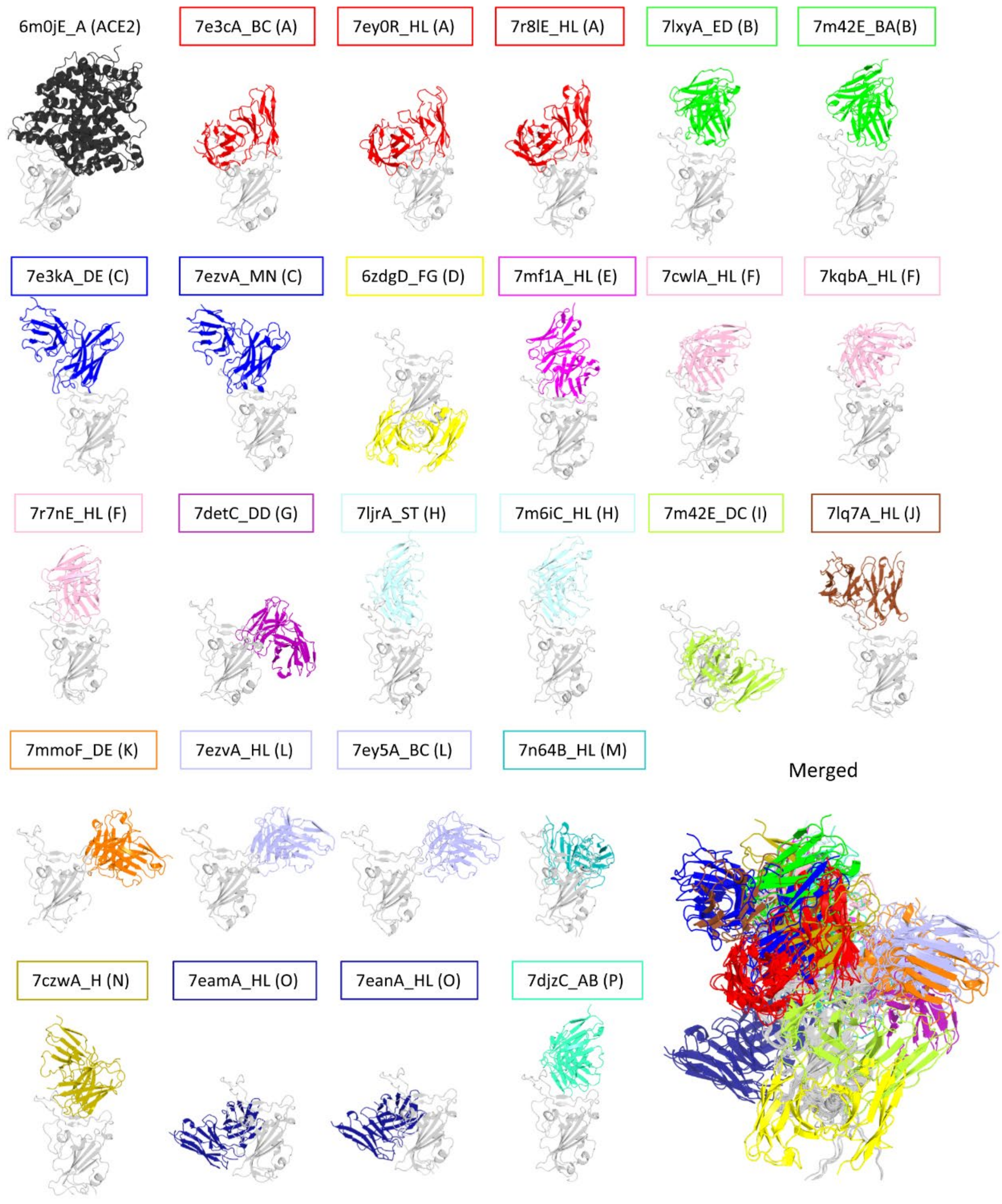


**Supplementary Figure 6. The visualization of 25 SARS-Cov-2 RBD-antibody complexes.** The epitope cluster is given followed by the query name. Each RBD in the complex was aligned as the orientation of the RBD in RBD-ACE2 binding pose.
